# Assessing Environmental RNAi in a Non-Model Organism

**DOI:** 10.1101/860338

**Authors:** Mosharrof Mondal, Jacob Peter, Obrie Scarbrough, Alex Flynt

## Abstract

RNA interference (RNAi) regulates gene expression in most multicellular organisms through binding of small RNA effectors to target transcripts. Exploiting this process is a popular strategy for genetic manipulation in invertebrates and has applications that includes control of pests. Successful RNAi technologies are dependent on delivery method. The most convenient method is likely feeding which is effective in some animals while others are insensitive. Thus, there is a need to develop RNAi technology on a per-species basis, which will require a comprehensive approach for assessing small RNA production from synthetic nucleic acids.

Using a biochemical and sequencing approaches we investigated the metabolism of ingested RNAs using the two-spotted spider mite, *Tetranychus urticae*, as a model for RNAi insensitivity. This chelicerae arthropod shows only modest response to oral RNAi and has biogenesis pathways distinct from model organisms. To identify RNAi substrates in *T. urticae* we characterized processing of synthetic RNAs and those derived from plant transcripts ingested during feeding. Through characterization of read size length and overlaps of small RNA reads, visualization methods were developed that facilitate distinguish *trans-*acting small RNAs from degradation fragments.

Using a strategy that delineates small RNA classes, we found a variety of RNA species are gated into spider mite RNAi pathways, however, potential mature trans-acting RNAs appear very unstable and rare. This suggests spider mite RNAi pathway products that originate as ingested materials may be preferentially metabolized instead of converted into regulators of gene expression. Spider mites infest a variety of plants, and it would be maladaptive to generate diverse gene regulators from dietary RNAs. This study provides a framework for assessing RNAi technology in organisms where genetic and biochemical tools are absent and benefit rationale design of RNAi triggers.

## BACKGROUND

RNAi is a mode of gene silencing where small 18-30 nucleotide (nt) RNAs associated with Argonaute (Ago) proteins bind to complementary transcripts^1^. In animals, small RNAs are separated into three major classes; microRNAs (miRNAs), small-interfering RNAs (siRNAs), and Piwi-interacting RNAs (piRNAs)^2^. While a role for miRNAs in gene-regulatory networks is highly conserved the function of siRNAs and piRNAs varies by clade. When RNAi is exploited to inhibit gene expression in invertebrates a common strategy is delivery of long dsRNA for processing into siRNA class small RNAs by the RNase III enzyme Dicer^3^. Separate from most other animals, arthropods have multiple Dicers, which at least in *D. melanogaster* segregates miRNA biogenesis from siRNAs^4^. A potential benefit of distinct miRNA/siRNA pathways could be to allow rapid evolution of siRNAs for antiviral or genome defense, something that is incongruent with conservation of miRNA function^5^.

A consequence of the rapidly evolving nature of siRNAs is inconsistency in RNAi efficacy across arthropods. For example, long dsRNA is effective in beetles, but not in moths and butterflies^6^. Exploiting sensitivity to dsRNA in beetles has already yielded commercial products to combat the coleopteran western corn root worm^7^. However, similar products have not been developed for other pests. This suggests that differences in RNAi response is a major complication that limits application of gene silencing technology. In such an application, animals will need to ingest molecules like long dsRNA, which adds further barriers such as destruction of molecules in gut lumen prior to entry into cells^8^. Moreover, preventing incidental entrance of exogenous RNA into RNAi pathways is probably a desirable trait, especially for polyphagous arthropods which are likely to ingest a great variety of RNA structures and sequences. Thus, methods to characterize metabolism of exogenous RNAs in target pests becomes valuable for identifying RNA species that will be strong candidates for technology development. In particular methods for non-model organisms that lack genetic tools and molecular biology reagents like antibodies for dissecting small RNA biogenesis would be beneficial.

The small RNA effectors of RNAi are particularly amenable to high-throughput sequencing approaches due to their short length. During library creation naturally occurring RNA ends and genome strand are preserved through sequential RNA ligation. By analyzing read sizes, nucleotide content, and overlaps between complementary, alternate-strand mapping reads; class and biogenesis patterns can be elucidated. In arthropods Dicer products (siRNAs and miRNAs) are typically 19-23 nt long. The RNase III domain of Dicer causes staggered cleavage of dsRNAs that leaves 2nt 3’ overhangs between strands of the siRNA duplex. In contrast, piRNAs are represented by longer 25-30nt reads. Biogenesis of piRNAs involves cleavage by piwi proteins guided by the activity of pre-existing piRNAs. Ago and Piwi proteins may exhibit slicer activity which cuts the base of target transcript 10 bases from the 5’ end of the bound small RNA. After piwi cleavage, the fragmented transcript is converted into a new piRNA directly or by alternative nucleases. Each of these biogenesis modes can be determined through sequencing data analysis independent of genetic tools.

To address this, we describe a comprehensive strategy for examining and visualizing exogenous RNAs that give rise to RNAi effectors. Our approach combines an isolation method alongside sequencing data analysis of small RNA sequencing libraries. To development this strategy we used the two-spotted spider mite. A variety of plants are infested by spider mites, and it is itself a prime candidate for new pest control technologies as it can rapidly develop resistance to conventional pesticides^9^. Several studies have tested *T. urticae* for sensitivity to dsRNA, finding saturating exposure like soaking in dsRNA solution is required for robust phenotypes^10,11^. Together this makes *T.urticae* an ideal pest to evaluate as it will reveal a profile of suboptimal environmental RNAi. Furthermore, mites are not insects but rather chelicerae arthropods, which harbor RNA-dependent RNA polymerases (Rdrps), a factor in nematodes and plants with a role in potentiating RNAi through amplifying dsRNA. However, there is little functional evidence that animal Rdrps, other than those found in nematodes, have a role in RNAi pathways that interact with exogenous RNAs^12,13^,^14^. Thus, this approach may also reveal a Rdrp signature for this group. Additionally, spider mites unlike *D. melanogaster* have a central role for siRNAs in genome surveillance^15,16^.

By applying this strategy, we find a variety of exogenous RNAs are processed into small RNA following ingestion by spider mites. Interestingly, the most active configuration is not the expected long dsRNA. We find a wide range of plant produced RNAs appear to be converted into small RNAs after ingestion. However, very few of the RNAs seem to associate with regulatory complexes, something that is affected by concentration of RNAs consumed. This suggests a scenario where foreign RNAs appear to transit the spider mite RNAi pathway but are diverted from regulatory complexes.

## RESULTS

Analysis of environmental RNAs competent to become *trans*-acting is complicated by the presence of fragments in samples. Immunopurification of Ago complexes followed by isolation and sequencing of associated RNAs can be used to validate small RNA identity^17^. However, validated antibodies necessary for this approach are unavailable for all but a handful of organisms. To address this problem, we used Hi Trap QFF chromatography, which can be used to enrich for Ago/Piwi-associated small RNAs (Fig 1)^18^. Unbound RNAs in animal lysates are retained on the on the resin while highly basic Ago/Piwi proteins pass through. When used on spider mites, a single clear peak of small RNAs is observed (Fig 1a). In comparison, total RNA samples show many additional RNAs (Fig 1b). After library creation and sequencing, size distribution of column purified RNAs shows a lower proportion of reads under 20 nt and an enrichment of piRNA sized (25-28nt) species (Fig 1c). We also observed three times the representation of miRNAs (Fig 1d). Using this purification strategy, we investigated the processing of exogenous dsRNAs into *bona fide trans-*acting small RNAs.

**Figure 1.**
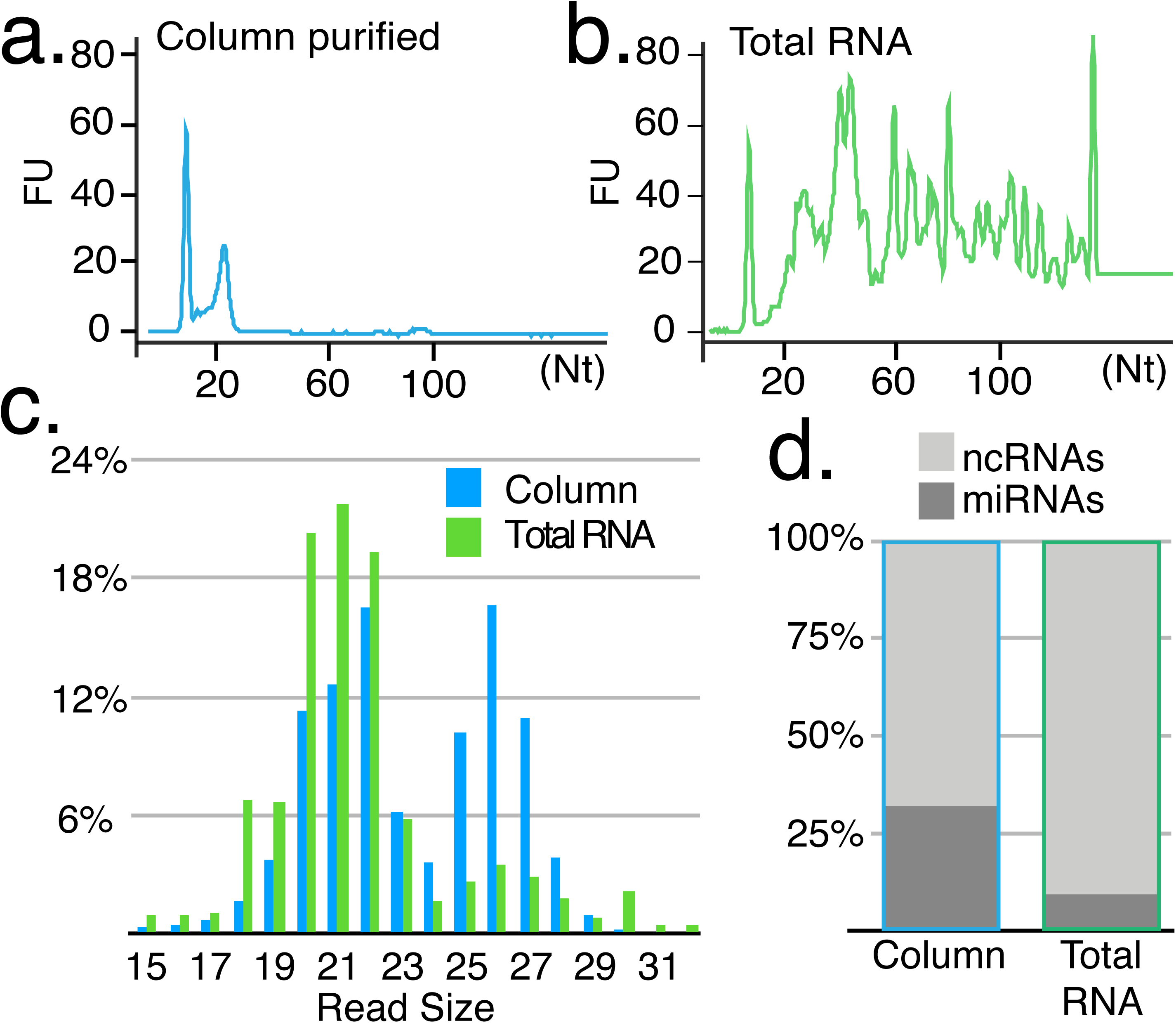
Enrichment of small RNAs with Hi Trap QFF columns. **a.** Bioanalyzer trace of RNA sizes following column purification compared to (**b**) RNAs extracted with TRI-reagent. **c.** Size distribution of reads from sequencing column purified or total RNA. **d.** Greater recovery of miRNAs (microRNAs), relative to ncRNA-derived transcripts.

Following previously reported methods, we sought to induce maximum gene silencing by soaking animals in dsRNA target to spider mite Actin and GFP (Fig 2 & Supp Fig 1). Following exposure to dsRNA, animals were processed into a lysate and small RNAs were isolated with the Hi Trap approach described above. Following sequencing, small RNA reads were aligned to actin and GFP sequences, which showed significant accumulation of 18-21 nt RNAs at the target region (Fig 2a & Sup Fig 1). For untreated conditions after Hi Trap enrichment or from total RNA the accumulation was absent with very few alignments in the enriched sample and many apparent degradation fragments for total RNA (Fig 2a). We also found actin and GFP siRNAs were not heritable as none were present in embryos sired by soaked mites (Sup Fig 1b,c).

**Figure 2.**
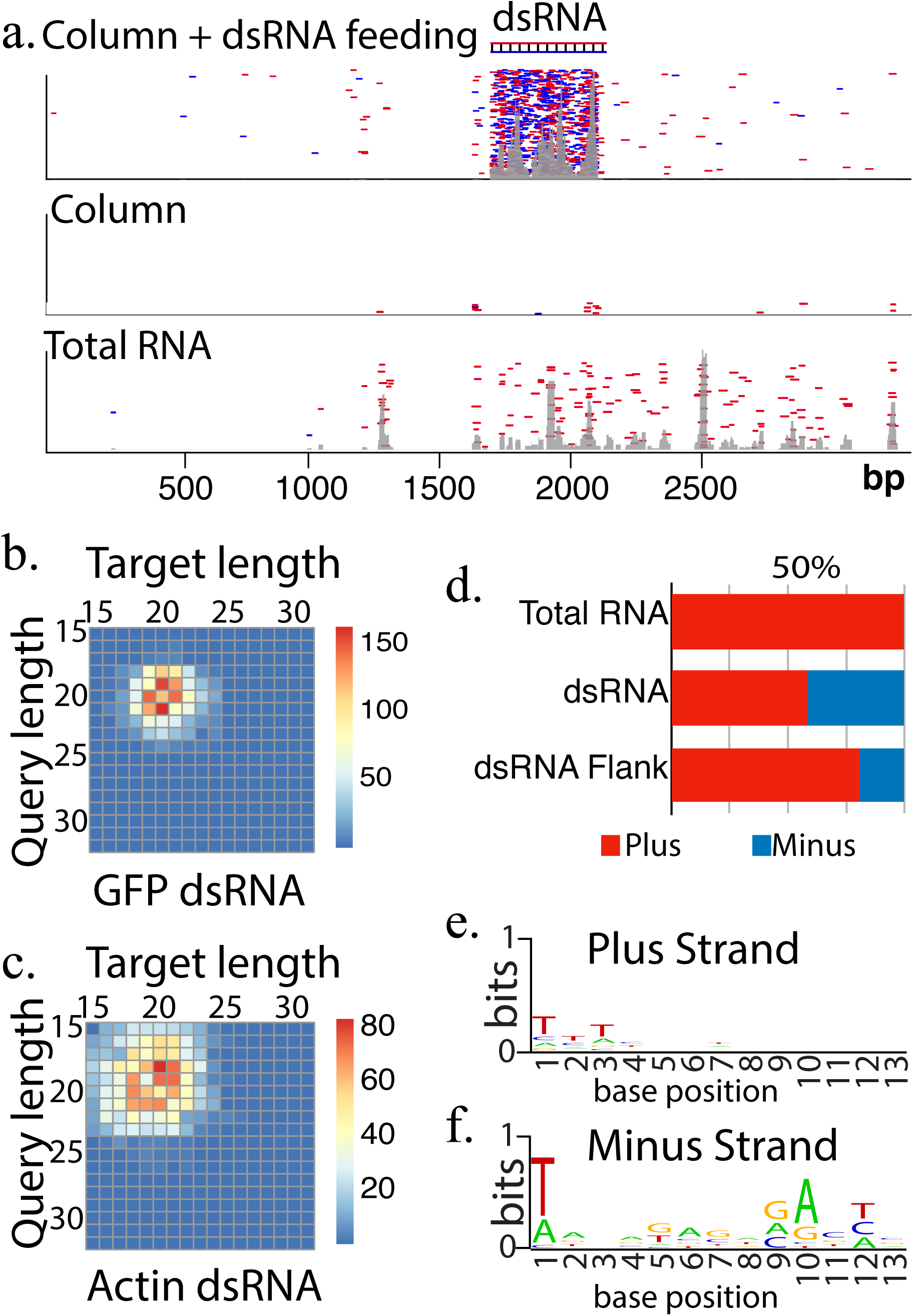
Fate of ingested synthetic long dsRNA. **a.** Alignment of reads to the Actin locus sequenced from column purified RNAs following feeding with dsRNA (top panel), and no feeding (middle and bottom panels; column and total RNA respectively). **b-c**. Number of overlap sequencing read pairs of different sizes derived from dsRNA targeted to GFP (**b**) or Actin (**c**). **d.** Portion of plus or minus strand mapping in Total RNA, dsRNA targeted region, or flanking region when dsRNA is fed. **e-f.** Seqlogo plot showing sequence bias for dsRNA flanking region mapped RNAs either to the plus (**e**) or minus strand (**f**) of Actin.

We then sought to verify Dicer processing of long dsRNA-derived siRNAs by examining overlaps between complementary pairs of small RNA reads^19^. Specifically, the abundance of small RNA pairs with 2nt overhangs, characteristic of RNase III cleavage, were summed and plotted in a matrix of pair lengths (Fig 2b,c). This reveals the sizes of small RNAs found in duplexes that have a signature of Dicer cleavage. In total ∼45% of Actin mapping reads showed signs of Dicer activity consistent with the centrality of this enzyme in the processing of dsRNA, and further validates the purification strategy for examining functional siRNAs. Common pairs were the expected 18-21 nt length and tended to be offset by 1-2 nts (Fig 2b,c). Actin siRNAs showed more diversity in pair lengths compared to GFP siRNAs, suggesting capture of target cleavage products by RNAi pathways occurs when a target transcript is present. We also observed many antisense reads mapping outside the target region that were not present in the total RNA library. This potentially could be the result of Rdrp activity producing antisense transcripts in response siRNA targeting (Fig 2d). We then inspected 5’ end sequence biases of the actin reads in the dsRNA flanking regions (Fig 2e,f). While there is little sequence similarity in sense mapping reads, antisense have clear 5’ terminal “T” and “A” 10 position. This is highly suggestive of piRNAs, which exhibit these features as a result of piRNA “ping-pong” processing^20^. In spider mites siRNAs and piRNAs cooperate for genome surveillance^15^. These results suggest this may extend to siRNAs derived from exogenous sources, which likewise appear to be able to trigger piRNA production.

Next, we investigated another source of exogenous RNAs–those acquired from plant hosts to assess the universe of RNA species that infiltrate RNAi pathways. Spider mites infest many plants but are readily reared on bean plants (*Phaseolus vulgaris*). In sRNA sequencing libraries created from total RNA extracted from bean-raised animals ∼3.13% map perfectly to bean sequences. Of these reads, 85% align exclusively while 15% also map to the mite genome. When Hi Trap purified libraries are aligned to plant sequences there is a 10-fold decrease in mapping rates, and a substantial shift in the proportion of reads that align to both mites and the plant (67%). Hi Trap mite purified RNA was also extracted from animals raised on *Arabidopsis thaliana*, which is a poor host^21^. We observe reduced plant aligning reads at a rate of 0.09%, consistent with reduced intake of plant materials. 57% of reads from this sample also mapped to the mite genome. Plant mapping RNAs derived from mites have features of mite small RNAs, such as a peak of read sizes expected for siRNAs (Sup Fig 2). Comparison to public small RNA sequencing libraries from the plants themselves showed shifts in the dominant read sizes. Thus, the RNAs in the mite library are unlikely mature plant sRNA contaminants.

Simultaneously, a substantial portion of the RNAs do appear to be generated from plant transcripts. The Dicer overlap pairs from mites derived RNAs that map to plant sequences exhibited a pattern similar to long dsRNA and the mite genome (Fig 2b,c,3a, Sup Fig 3a). The same pattern was not seen with total RNA library mapping to plant (Fig 3a). Endogenous plant sRNAs were also different, showing the greatest number of pairs at 24 nt, which is consistent with the 24 nt Dicer products found in plants (Fig 3a, Sup Fig 3c)^22^. Strikingly, the plant mapping reads from both Hi Trap and total RNA samples do not recapitulate this pattern when aligned to the mite genome (Fig 3a). Thus, it would seem the plant mapping RNAs derived from mite samples are for a large part produced from plant transcripts despite co-mapping to the *T. urticae* genome. Interestingly, plant mapping reads that do map back to the spider mite genome overlap with longer-sized reads in mites, again potentially tying piRNAs to siRNAs in this animal.

**Figure 3.**
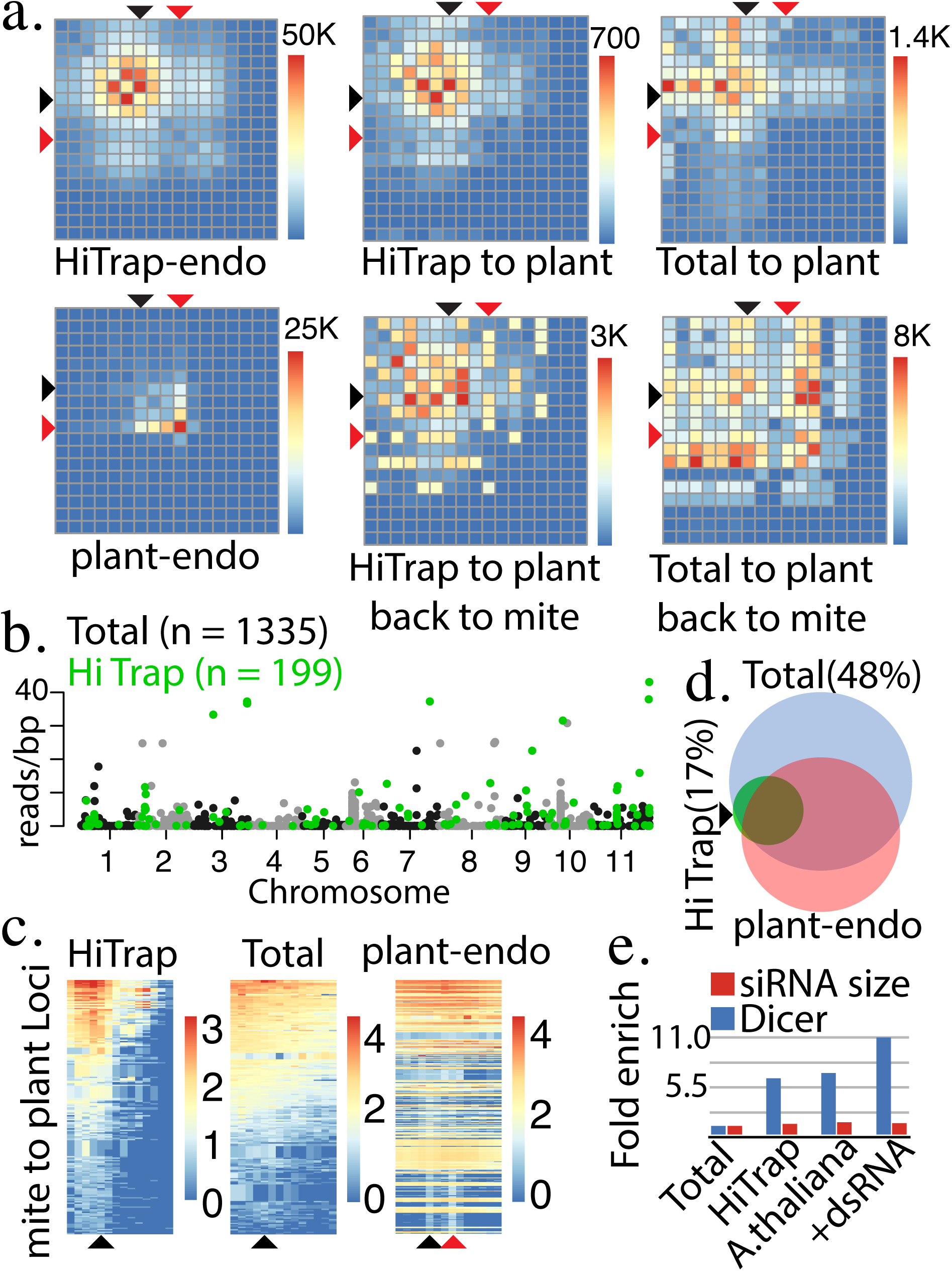
Metabolism of plant RNAs and identities of transcripts that enter small RNA pathways. **a.** Number of Dicer overlap read pairs for HiTrap purified mite derived RNAs mapped to the *T. urticae* genome (top left) and those that map to P. vulgaris sequences (top mid). Dicer overlap reads plotted for reads cloned from total RNA (top right, from endogenous plant RNAs (bottom left), and plant mapping RNAs from HiTrap purified (bottom mid) and total RNA samples mapped back to mite (bottom right). Black arrow shows 21nt size RNAs, red arrow 24nt **b.** 1335 Loci in *Phaseolus vulgaris* (bean) with at least two read depth and greater than 40 nucleotides long identified using RNAs derived from mites. Loci are plotted by chromosome and number of reads per base pair. Green labeled loci are the 199 subset found in HiTrap dataset. **c.** Heatmaps showing size distribution of small RNA reads for loci identified in (**b**), using mite-derived HiTrap RNAs (HiTrap), total RNA extracted (Total), and bean tissue derived (endo-sRNA) libraries. Black arrow shows 21nt size RNAs, red arrow 24nt. **d.** Venn diagram showing overlap between loci plotted in (**b**) and loci expressing endogenous small RNAs. Percentage indicates portion of mite library-identified loci that don’t overlap with endogenous sources of small RNAs. **e.** Enrichment of plant mapping reads that exhibit Dicer overhangs in HiTrap purified samples from P. vulgaris, A. thaliana, and mites fed dsRNA raised on P.vulgaris. The portion of reads between 19-23nt long (siRNA size) also plotted.

We then annotated loci that potentially transmit RNAs to mites based on read alignment density to plant genomes (Fig 3b, Sup Fig 3d). Using the total RNA dataset we uncovered 1335 loci and 199 with the Hi Trap sample. Nearly all the Hi Trap loci overlapped with the ones annotated with total RNA, however, this was not tied to relative expression level. Examining the size distribution of reads mapping for each Hi Trap annotated locus showed bias similar to synthetic dsRNAs (Fig 3c, Sup Fig 1a, Sup Fig 2c,d). The pattern was not seen for these loci when using total RNA library alignments or a plant endogenous small RNA library (Fig 3c). Next, we intersected endogenous plant small RNAs loci with those called from mite derived RNAs libraries (Fig 3d). Generally, there was significant overlap with Hi Trap libraries but not total RNA (Fig 3d). However, the Hi Trap RNAs are not generated from well-processed plant small RNAs like miRNAs, which were only found in the total RNA dataset. This suggests that while plant RNAs that have features of small RNAs precursors are more likely to enter mite RNAi pathways, it is probably the precursor form of the RNA that is taken up. We also observe enrichment of Dicer overlap reads as a portion of mapping reads in Hi Trap samples with an apparent further enrichment for animals exposed to the synthetic dsRNA (Fig 3e). Thus, it would appear super abundance of dsRNAs in diet could saturate turnover enzymes and leads to greater stability of the small RNAs derived from dietary sources.

To explore the greater accumulation of Dicer products after dsRNA feeding, we focused specifically on alignments of mite derived libraries to plastids as these organelles do not have Dicer activity. Potential dsRNA/shRNA substrates are not processed prior to ingestion^23^. Plastids also lack well defined transcriptional units, leading to potential widespread formation of dsRNA^24^. Consistent with precursors of small RNAs being transmitted we observe over-representation of chloroplast sequences in all mite-derived libraries (Fig 4a). Greater accumulation was seen in bean fed mites versus *A. thaliana* where less plant material is ingested. Interestingly, we also observe dsRNA feeding correlated with the fold increase in plastid genome coverage. This appears to be in part caused by a shift to dsRNA substrates. The amount of genome coverage for single-stranded mapping is similar between dsRNA fed and non-fed mites while the portion covered by dual strand mapping nearly doubles in the fed animals (Fig 4b).

**Figure 4.**
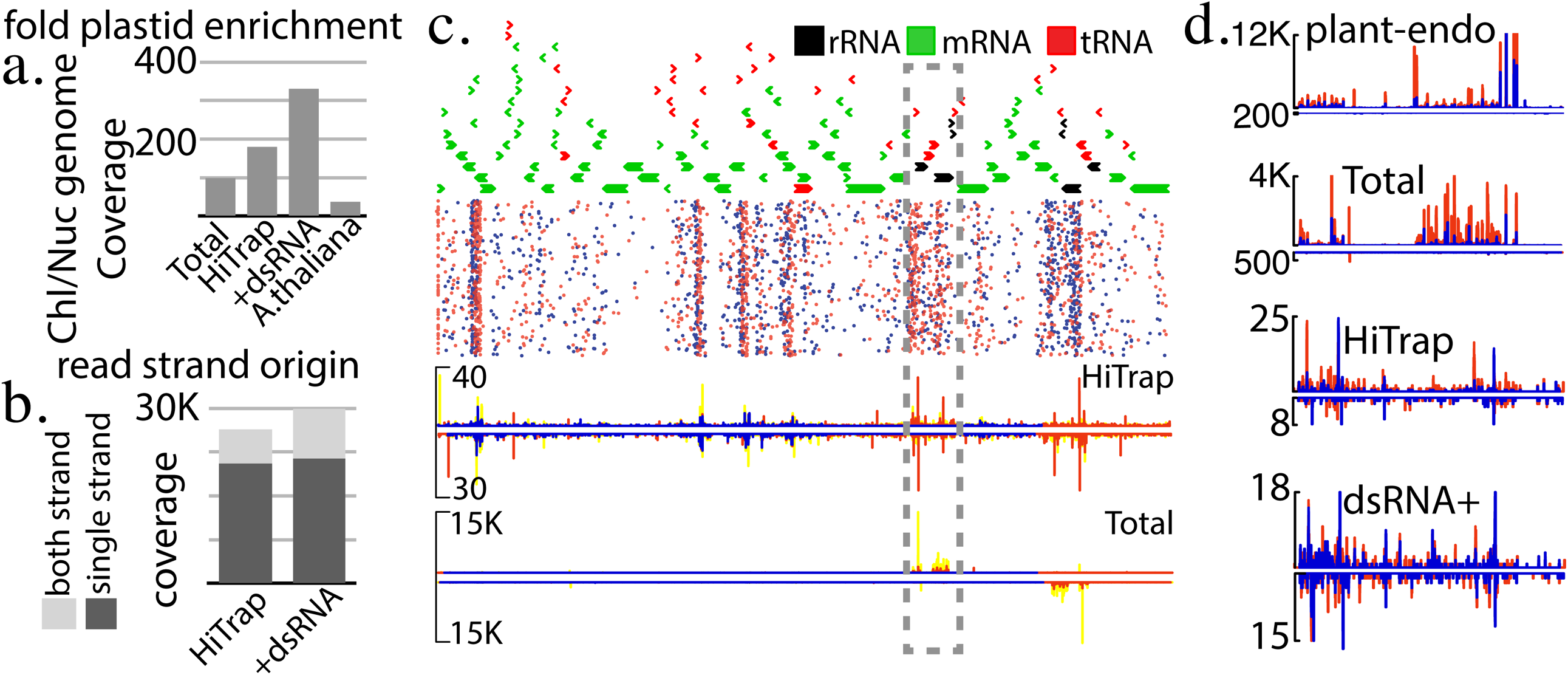
Chloroplast mapping RNAs. **a.** Coverage of chloroplast relative to nuclear genome for RNAs recovered from total and HiTrap purified RNAs from mites raised on bean plants, bean plants after dsRNA feeding, and *A.thaliana*. **b.** Plastid genome coverage simultaneously on both strands or single strand. **c.** Distribution of reads mapping to *P.vulgaris* chloroplast genome. Top trace shows chloroplast genes. Next panel, alignment of individual reads from HiTrap purified samples. Bottom two panels show read densities for HiTrap and total RNA samples. Orange represents 19-23nt reads that map in multiple positions, and blue unique mapping positions for the 19-23nt size range. Yellow shows densities of all size ranges (15-30nt) **d**. Density plots showing read accumulation in the region in (c) highlighted by the gray dashed box. Library used to generate plot above, mapping separated by strand. Red represents all read ranges (15-30nt), and blue siRNA size only (19-23nt). Y-axis is maximum density.

Generally, small RNAs map to regions of high gene density and rRNA loci (Fig 4c). In total RNA libraries the majority of reads map to a single strand of the rRNA loci. In contrast, the Hi Trap dataset show more consistent dual strand mapping, even at rRNA, albeit at a greatly reduced rate (Fig 4d). In animals fed dsRNA, dual-strand coverage is even more equal with a greater portion of siRNA-sized (19-23nt) transcripts. This suggests that among plastid RNAs ingested by mites a small, yet clear subset enter RNAi pathway. Prior exposure to dsRNA appears to stabilize the on-pathway RNAs which we propose is the result of saturating processing machinery. Thus, it would appear that siRNAs generated from exogenous RNAs are very unstable being preferentially turned over rather than incorporated into regulatory complexes.

Based on our finding that structured plant derived RNAs enter spider mite RNAi pathways we investigated the activity of short hairpin RNAs that mimic distinct, known miRNA-type biogenesis (Fig 5a). The ability of four different configurations of synthetic hairpin RNAs to trigger silencing of Actin was tested alongside long dsRNA also targeted to Actin by soaking in *in vitro* synthesized molecules. qPCR was used to quantify target knockdown (Fig 5b). The short hairpins were designed to transit: canonical miRNA biogenesis (shRNA), Ago processed short loop RNAs that mimic miR-451 with and without a G-C clamp at the hairpin base (SL1 & SL2), and a G-C clamp stabilized RNA with a loop sequence complementary to actin (Loop)^25,26^. GC clamps were added to the structure bases of the RNAs to increase stability as this high energy fold is a challenging substrate for RNases^27^. After soaking, relative abundance of actin transcripts was assessed showing the reported ∼40% reduction for long dsRNA^10^. Three of the structured RNAs showed a greater degree of knockdown with a reduction of transcript accumulation reaching nearly 60%, suggesting that the siRNA pathway of *T. urticae* may not be the optimal mode of RNAi to exploit for gene silencing (Fig 5b).

**Figure 5.**
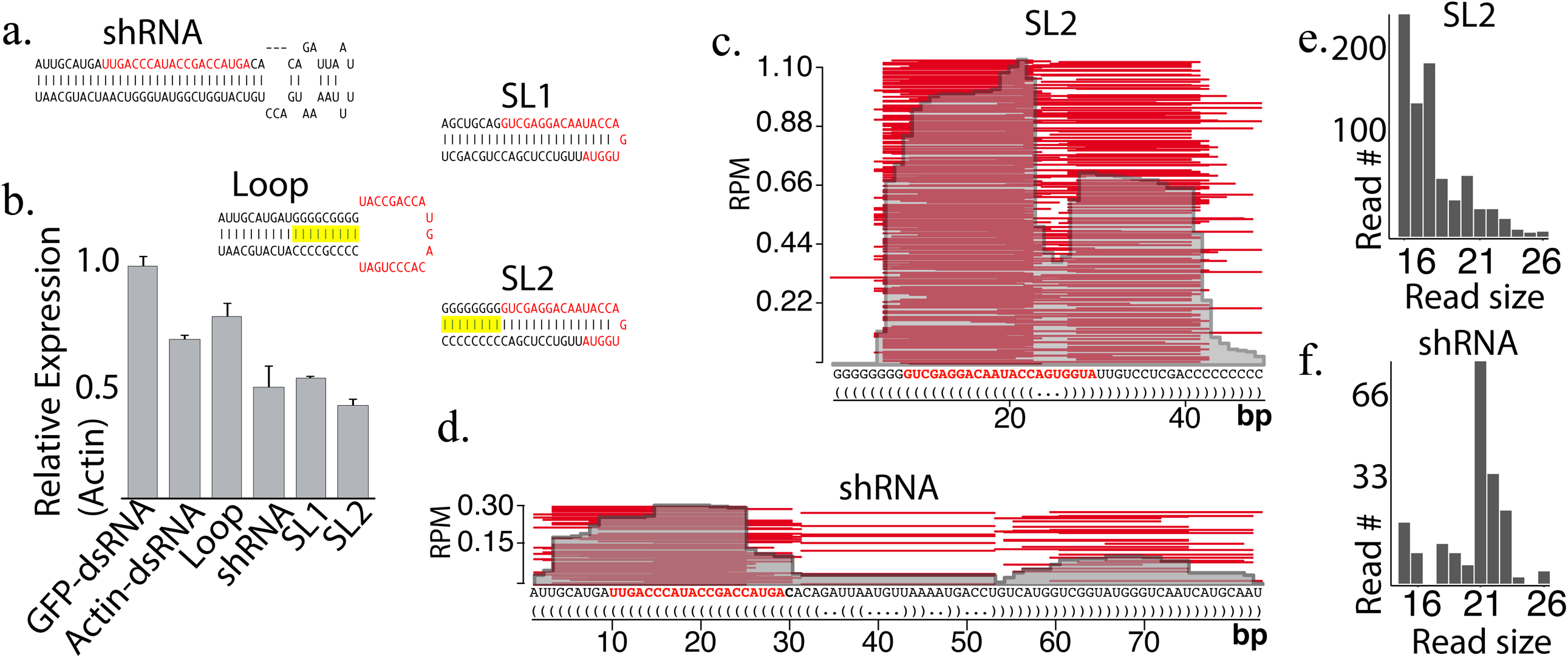
Fate of synthetic short hairpin RNAs fed to spider mites. **a.** Actin targeted structured short hairpin RNAs fed to animals via soaking. Designs mimic modes of miRNA biogenesis: Dicer processed (shRNA), Ago processed short loop RNAs (SL1, SL2), and loop derived RNAs (Loop). Red sequence indicates portion targeted to Actin. Yellow highlight show G/C “clamp”. **b.** Percent expression of target gene (Actin) following ingestion of long double-stranded RNA (dsRNA) or short RNAs (**a**). Error bars are standard deviation. **c-d.** Alignment of RNAs sequenced from animals fed SL2 (**c**) or shRNA (**d**) synthetic RNAs. Red sequence shows portion targeted to Actin. **e-f.** Absolute number of reads mapping to short hairpin RNAs SL2 (**e**) and shRNA (**f**)

We also sequenced samples after feeding with the shRNA (Fig 5c) and SL2 (Fig 5d) RNAs. SL2 derived small RNAs were more abundant but showed smaller fragments than shRNA, consistent with being an atypical RNAi substrate (Fig 5e,f). For both RNAs, however, cleavage sites were non-uniform, and for SL2 there was little evidence slicer-mediated precursor processing was occurring as seen with miR-451, but rather standard dicer processing, suggesting that additional engineering could yield an even more superior gene silencing molecule. Nevertheless, many reads were complementary to Actin that were able to trigger knockdown. This together with the appearance of long dsRNA-derived siRNAs shows that a variety of ingested RNAs enter spider mite RNAi pathways, and that the designated siRNA/long dsRNA pathway may not be the most potent option for eliciting knockdown.

## DISCUSSION

Using the strategy described here we were able to provide a comprehensive view of environmental RNA processing in the two-spotted spider mite. Several novel aspects of small RNA biogenesis emerged that are not found in model organisms to include involvement of ping pong piRNA-like processing of transcripts targeted by dsRNA. Critically, these insights were possible without genetic manipulation or well-developed molecular biology reagents. An identical analysis could be performed on nearly any animal. The value of this approach is evident from the finding that many types of single-strand structured RNAs are processed by Dicer into small RNAs, which led us to identify several short hairpin RNAs that have greater gene silencing efficacy compared to long dsRNA.

Our analysis also has important implications for RNAi strategies. While we identified a wide range of ingested RNAs can access mite RNAi pathways, phenotypes are hard to induce in these animals with RNAi. We propose that this apparent contradiction may indicate that RNAi pathways in mites have another function: metabolism of double-stranded or structured RNAs. Identity of transcripts converted into small RNAs by mites is highly dynamic and tied to feeding state. Moreover, for an animal with hundreds of plant hosts, routine gating of ingested RNAs into multi-target gene regulators would be maladaptive. Preferential destruction of small RNAs based on origin is seen in *Drosophila*^*28,29*^. The mirtron subclass of miRNAs as well as siRNAs derived from latent viruses are poorly recruited to Ago complexes^28,29^. Navigating such a fate for small RNAs will be critical to effective RNAi technology. Avoiding the conversion of ingested RNAs into regulators could be beneficial to variety of animals that share RNAi factors and ecology with spider mites.

The primary value of the approach we describe here is that it is species agnostic. There are many species that would be prime candidates for new pest control technologies where RNAi efficacy is unclear. By evaluating a range of substrates both synthetic and dietary using our strategy better approaches could be developed. The synthetic hairpin RNAs that showed greater activity in spider mites are likely processed via miRNA biogenesis a much more conserved mechanism that arthropod siRNAs. A similar type of RNAi trigger may be a better choice for animals that have apparent resistance to long dsRNA. The biochemical and computational approach here would confirm this is situation in one of these species.

## CONCLUSIONS

This study describes a strategy for assessing small RNA biogenesis in non-model organisms. By pairing a species agnostic biochemical method with sequencing analysis appropriate for small RNAs, deep insights are possible. Our investigation depended on several distinct analyses that identified the class of the small RNA based on apparent processing. By comparing read lengths, overlaps with complementary reads RNAs could be identified as either piRNA, siRNA, or miRNA. Moreover, by combining Ago enrichment with small RNA sequencing dietary RNAs competent for RNAi can be characterized. This can lead to a better-informed design of RNAi triggers such as the short hairpin RNA found in this study.

## METHODS

### Mite handling

Mites were reared on two plants: *Phaseolus vulgaris* or *Arabidopsis thaliana.* Mites were removed from plant leaves by gentle tapping and collected into a microfuge tube. Both juveniles and adults were kept. Mite soaking was done by adding 200 *µ*L of a 160 ng/*µ*L RNA solution to collected animals. Soaking lasted overnight (15-16 hours) followed by rinsing in PBS (pH 7.4), and 0.1% Tween 20 solution. Afterwards, mites were collected on a paper towel and allowed to dry before being placed onto a *P. vulgaris* leaf inside of a large petri dish at room temperature. After five days, mites were collected, flash frozen and stored at −80°C.

### RNA synthesis

For the initial experiment, two dsRNAs were generated from cloned fragments of *T. urticae* actin and of Green Fluorescent Protein (GFP) from *Aequorea victoria* (accession numbers CAEY01002033.1 and FJ172221.1, respectively). Actin long dsRNA and GFP long dsRNA, were both approximately 350 nt long and created using the MEGAscript™ *in vitro* transcription kit (Ambion). Templates for dsRNA synthesis were PCR products amplified with primers encoding T7 RNA polymerase promoter sequences on their 5’ ends. Following *in vitro* transcription, lithium chloride precipitation was performed to purify products, and the resulting RNAs were annealed through heating and gradual cooling.

Four synthetic structures were designed based on RNAs shown to enter miRNA pathways. Templates for these structures were created by annealing long, ∼100 nt, oligonucleotides (supplement). The resultant dsDNAs encoded a T7 promoter at the 5’ end followed by the structured RNA sequence. RNAs were synthesized using the MEGAshortscript™ *in vitro* transcription kit (Ambion). The synthesis products were purified by phenol chloroform extraction, and ethanol precipitation.

### HiTrap Q FF column enrichment

Column enrichment of small RNA containing complexes used 1 mL HiTrap Q FF columns (GE Lifesciences) as previously described^18^. Briefly, columns were equilibrated following manufacturer instructions. First, 5 mL of start buffer (20 mM HEPES-KOH, pH 7.9) was applied and passed through the column (1 mL/minute). Then 5 mL of elution buffer (20 mM HEPES-KOH, pH 7.9, 1 M NaCl), and 10 mL start buffer (20 mM HEPES-KOH, pH 7.9, 100 mM KOAc) were applied sequentially. Collected animals were washed several times with PBS, pH 7.4, followed by flash freezing and grinding with a mortar and pestle. 1 mL chilled binding buffer (20 mM HEPES-KOH pH 7.9, 100 mM KOAc, 0.2 mM EDTA, 1.5 mL MgCl_2_, 10% glycerol, 0.2% PMSF, 1 mM DTT, 1X Roche EDTA-free protease inhibitor cocktail) was mixed with the pulverized animals. The lysate was then clarified by centrifugation. Cleared lysate was applied to the column and passed at a speed of 1 mL/minute. The column was washed with binding buffer followed by elution buffer (binding buffer with 300 mM KOAc). An equal volume of acid phenol-chloroform was added to the tube, followed by rocking at room temperature. Following phase separation, RNAs were isopropanol precipitated and resuspended in 30 *µ*L of ddH_2_O. Total RNAs extracted from mites used TRI reagent and followed manufacturer protocols.

### RT-qPCR

Total RNA from spider mite samples was used for cDNA synthesis (Thermo Fisher T-7 kit) using random hexamer primers. cDNAs were used in qPCR assays with SYBR Green real-time PCR master mix (Thermo Fisher), following manufacturer protocols. In the qPCR assay, actin transcripts were normalized by assessing levels of 18S ribosomal RNA. Primer sequences for actin and were previously published^10^.

### RNA Sequencing and Analysis

RNA obtained from both column extraction and total RNA extraction were sequenced from an Illumina TrueSeq Small RNA Library Prep Kit library. Sequencing occurred on an Illumina Nextseq500 or Miseq machine using a single-read 50 base pair protocol.

Small RNA TruSeq libraries were initially processed using Fastx toolkit to remove adapter sequences^30^. Libraries were normalized for number of reads by subsampling with the Seqtk program. For experiments looking at dsRNA processing 50M reads were used, when comparison of plant derived RNAs 14M were used. Mapping of reads to mite and plant genomes used the Bowtie program with -a -v0 -m200 parameters to find all perfect alignments while also excluding extremely low complexity sequences^31^. Mapping exclusively to actin and GFP sequences used -v0 for only perfect matching RNAs, and chloroplast alignments used -v 0 -a to uncover all perfect alignments.

Alignments were then processed with Samtools and Bedtools to find regions of expression based on read density and merging of juxtaposed features^32,33^. Bedtools was also used to quantify alignments per feature and determine the intersection of genomic loci with library alignments. All mite and plant sequences as well as their annotations were taken from public databases^16,34-36^. Size distributions were calculated from alignments converted back to “fastq” format, which were then parsed to determine number of reads corresponding to lengths between 15-31 nt long.

A python-based algorithm was used to find overlapping read pairs that represent Dicer cleavage produced small RNA duplexes^37^. Small RNAs of 15-31 nts were used to query target pairs, also 15-31 nts long, that exhibited an overlap of 2 nts less than the size of query RNA. The number of reads for each pair was plotted to show a heatmap matrix. Visualizations were created using the R packages: ggplot2, qqman, Pheatmap, and Sushi packages^38-41^. All mite sequencing data can be accessed for the BioProject ID: PRJNA591169. Public libraries used for plant mapping are: SRR7738374 (*P.vulgaris*), and SRR7947145 (*A.thaliana*)

## ACKNOWLEDGEMENT

The work was supported from NSF-MCB (Award ID: 1616725), Mississippi INBRE program (P204M103476) from the National Institute of General Medical Science, and the UMMC Molecular and Genomics Facility which is supported, in part, by funds from NIGMS, including Mississippi INBRE, Obesity, Cardiorenal and Metabolic Diseases-COBRE (P20GM104357), and Mississippi Center of Excellence in Perinatal Research (MS-CEPR)-COBRE (P20GM121334).

We would like to thank Eric Riddick and Guadalupe Rojas from the USDA ARS center in Stoneville MS for supplying founders for a *Tetranychus urticae Koch* colony.

