## supplemental_Figures for "Assessing Environmental RNAi in a Non-Model Organism"

### Supplemental Information:

#### Oligos to clone short RNAs

SL1s: TAATACGACTCACTATAGAGCTGCAGGTCGAGGACAATACCAGTGGTATTGTCCTCGACCTGCAGCT
SL1as: AGCTGCAGGTCGAGGACAATACCACTGGTATTGTCCTCGACCTGCAGCTCTATAGTGAGTCGTATTA

SL2s: TAATACGACTCACTATAGGGGGGGGGGTCGAGGACAATACCAGTGGTATTGTCCTCGACCCCCCCCC
SL2as: GGGGGGGGTCGAGGACAATACCACTGGTATTGTCCTCGACCCCCCCCCCTATAGTGAGTCGTATTA

loopS: TAATACGACTCACTATAGGGGCGGGGTACCGACCATGACACCCTGATCCCCGCCCC
loopAS: GGGGCGGGGATCAGGGTGTCATGGTCGGTACCCCGCCCCTATAGTGAGTCGTATTA

shRNAs:
TAATACGACTCACTATAGTGCATGATTGACCCATACCGACCATGACATGTTTAGTTATTGTCATGGTGGCTATGGGTCAATCATGCAC
shRNAas:
GTGCATGATTGACCCATAGCCACCATGACAATAACTAAACATGTCATGGTCGGTATGGGTCAATCATGCACTATAGTGAGTCGTATTA

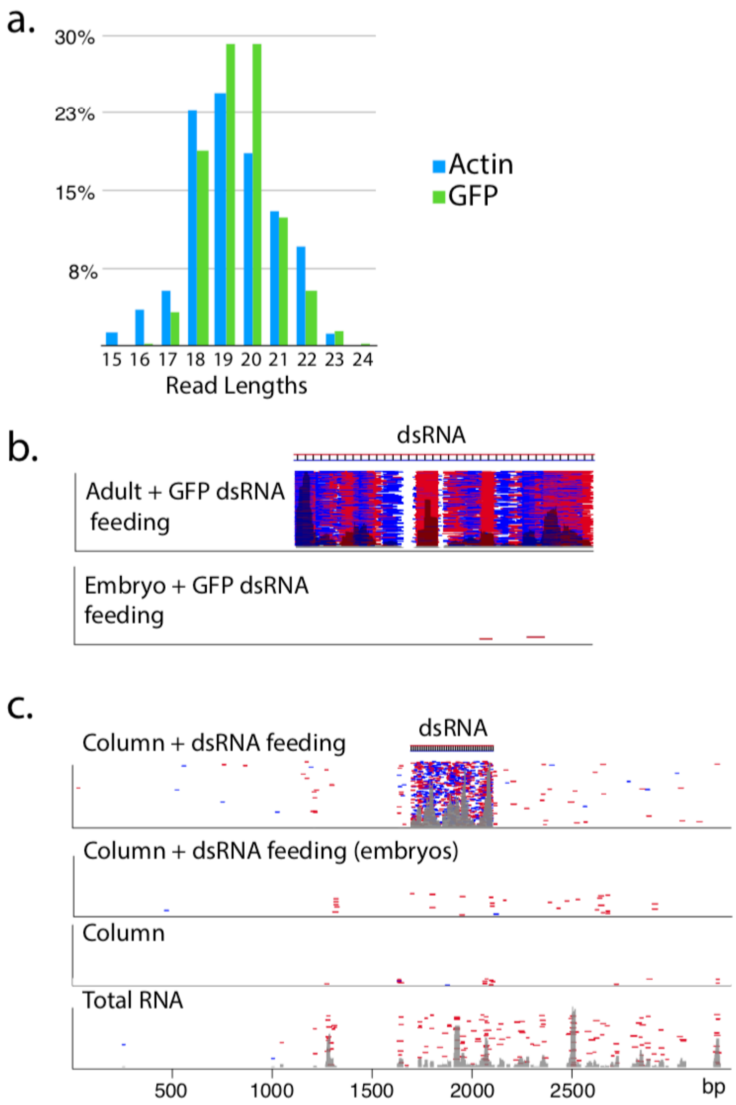


**Supplement Figure 1. Size distribution and mapping of dsRNA derived reads. a.** Percentage of reads at length 15-30nt mapping to Actin (blue) and GFP (green) **b-c.** dsRNA Alignment of reads to GFP(**b**) and Actin(**c**) following feeding of cognate dsRNA from Hi trap purified RNAs. For both targets, accumulation of RNAs in adults is shown as well as their progeny, showing little transmission of RNAs to the next generation after feeding of dsRNA. For actin, traces are shown for adults not fed dsRNA, and from a library constructed from total RNA.

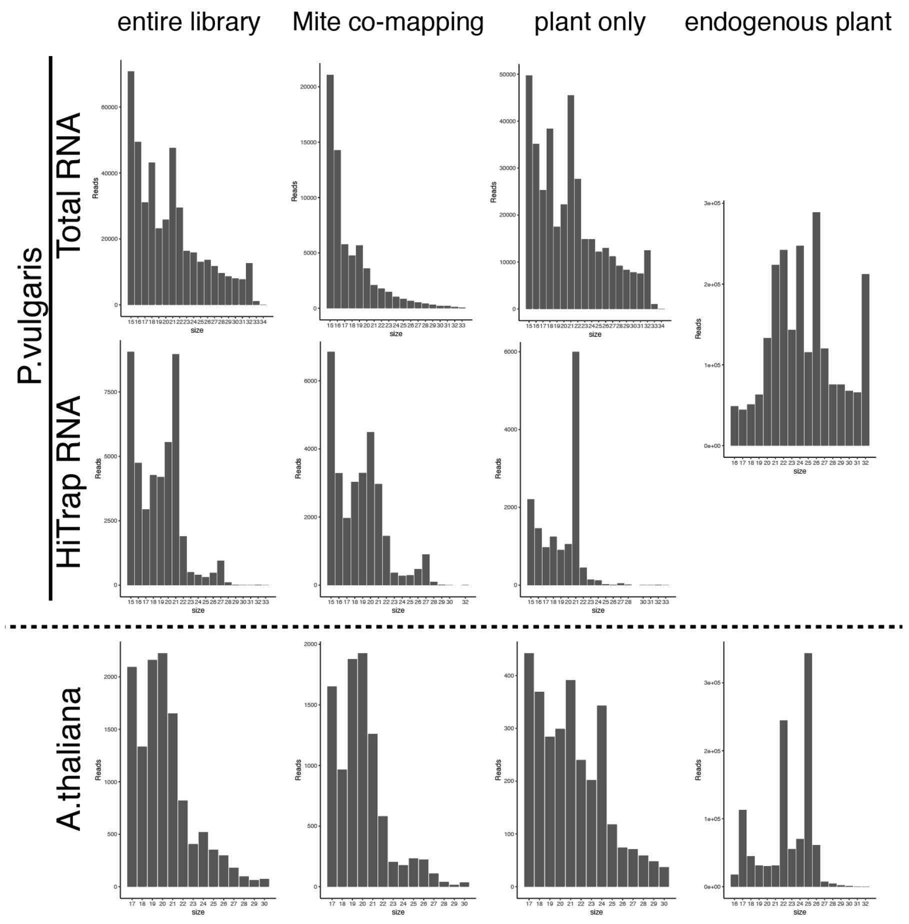


**Supplement Figure 2. Size distribution of plant mapping RNAs derived from mites.** Size of Reads on x-axis, number of reads on y-axis. Left most graphs show mapping to indicated plant and sample type. The next column is reads that map to plants and mites. Third column is mite-derived reads that map to plants that do not also map to mites. On right is size distribution of small RNA libraries generated from plant tissue.

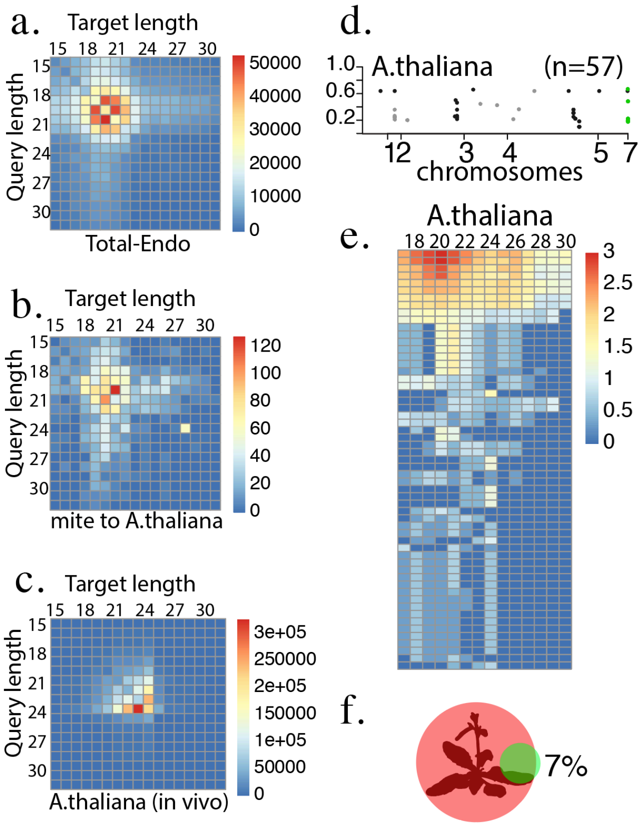


**Supplement Figure 3.** Number of overlapping read pairs of indicated size ranges for total RNA from bean fed mites mapped to the mite genome (a), from mites fed on *A.thaliana*, (**a**), and (**c**) read pair overlaps for *A. thaliana* endogenous (*in vivo*) plant RNAs. **d.** Loci in *A.thaliana* with a two read depth and greater than 40 nucleotides long. Loci are plotted by chromosome and bias towards siRNA-like (20-23nt) reads. Green labeled loci are those found in chloroplast. **e.** Heatmap showing size distribution of reads mapped to the loci identified in *A.thaliana.* **f.** Venn Diagram showing overlap of mite derived reads and endogenous siRNA producing loci in the plant

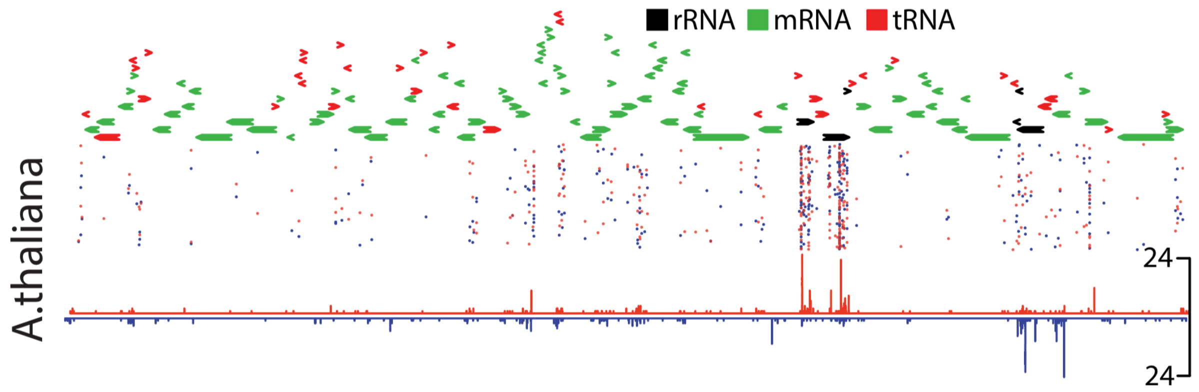


**Supplemental Figure 4.** Chloroplast mapping RNAs. Distribution of reads mapping to *A.thaliana* chloroplast genome. Top trace shows chloroplast gene. Mid panel, alignment of individual reads, and bottom a histogram of alignment densities.
